# *In vitro*, Evaluation for Consumption Ratio for certain plants and Toxicity and Biochemical Parameters of some Insecticides on *Eobania vermiculata*

**DOI:** 10.1101/2023.12.13.571426

**Authors:** A.S. Bashandy

## Abstract

Land snails are one of the most harmful agricultural pests to crops, causing economic devastation to a variety of plants. The purpose of this study is to determine the optimal food preference of land snails, *Eobania vermiculata*, and evaluate the mortality and biochemical activity of five pesticides using a leaf dipping technique against them in vitro. The findings revealed that in “no-choice feeding,” it was observed that land snails consumed a significantly higher amount of Cos lettuce leaves compared to leaves from other plants. However, chicory leaves were found to be the least preferred by the snails. On the other hand, “free choice feeding,” revealed that land snails showed a preference for Cos lettuce and cabbage leaves, which were consumed more frequently. Conversely, Komatsuna leaves had the lowest acceptance among the snails. Also, the results indicated the mortality rate increased with an increased concentration during exposure time. After three days, the lower concentrations of Sulfoxaflor and Fipronil caused a high mortality percentage in the animals compared with other pesticides, respectively. Moreover, after one month of exposure to 1000 ppm/100 ml, Chlorantraniliprole caused 46.43% mortality. But Sulfoxaflor and Fipronil exhibited (42.86%) mortality. They achieved LC_50_ (1010.48, 2501.93, and 1444.66 ppm/100ml), against tested snails, respectively. Nevertheless, Spinetoram and Spirotetramat caused a lower mortality rate. During seven days of exposure to LC_25_ for five pesticides by leaf dipping on E. vermiculata. They impacted some of the biochemical activity of two enzymes: alanine amino transaminase (ALT), aspartate amino transaminase (AST), total proteins (LP), and the lipid profile of E. vermiculata. The results showed that the activities of ALT, AST, and Triglycerides rose by 50%, with 108 u/l, 407 u/l, and 2064 mg/dl, respectively. Compared to the control and other compounds, Spirotetramat raised total cholesterol by 33 mg/dl. ALT and AST activity and Triglycerides lowered following Spinetoram and Fipronil treatment at 13 u/l, 32u/l, and (4 mg/dl) of TL, respectively. However, no substantial treatments or controls that influence TP, (Total cholesterol and LDH) or LP levels after the exposure period are available. Ultimately, the results indicated that both Cos lettuce and cabbage leaves were the preferred food choices for land snails, as observed in both feeding methods. However, it is important to note that these findings also highlight the need to consider the impact of insecticides on land snails. Incorporating these insecticides into a comprehensive management strategy to mitigate any negative effects of land snails while ensuring the overall well-being of the environment.

## Introduction

Mollusks are the animal kingdom’s second biggest phylum. It is classified into six taxonomic groups, with Class gastropods containing 80% of the Molluscan species. Gastropods are invertebrates and hermaphrodites with spirally coiled bodies (**Srivastava, 1992; Zala, *et al*., 2018**). Land mollusks are extremely damaging pests of vegetables, field crops, ornamental plants, fruit trees, and ecosystems (**Godan, 1983; Carlsson, *et al*., 2004**). Several species of snails and slugs are considered pests in agroecosystems around the world due to their high reproductive potential, nocturnal activity, and feeding behaviors, which cause crop damage and economic loss (**Routray and De, 2016; Ali, 2017; Das, *et al*., 2020**).

According to **Ismail (1997), 2004; Bayoumi *et al*., (2023)**, *Eobania vermiculata* species (Mollusca-Helicidae) are one of the most dangerous pests of numerous crops, vegetables, orchards, and ornamental plants.

The most convenient approach for managing terrestrial gastropods is the use of synthetic pesticides or particular molluscicides (**Radwan *et al*., 1992; El-Wakil and Attia, 1999; Moran *et al*., 2004; El-Shahaat *et al*., 2005; El-Shahaat *et al*., 2009; Eshr, 2014**).

Also, several efforts have been undertaken to identify molluscicidal effects using standard molluscicides such as metaldehyde and methiocarb, which are manufactured as baits (**Miller *et al*., 1988; Radwan, 1993**). Unfortunately, due to the poisonous nature of these compounds, they cause toxicity issues in non-target creatures as well as negative long-term consequences on the environment (**Homeida and Cooke, 1982; Smith *et al*., 1988**). Safe pesticides or molluscicides with unique mechanisms of action are so necessary.

Spirotetramat **(Movento 10% SC**), a spirocyclic tetramic acid derivative (Bretschneider *et al*., 2007), is a fully systemic insecticide for sucking pests. Spirotetramat is a lipid biosynthesis inhibitor (Nauen *et al*., 2006). The mode of action spirotetramat reduces fecundity and fertility and insect populations on all stages of pests.

Sulfoxaflor **(Transform 50%WG**) was approved as an insecticide in 2010, and it was used to protect diverse crops from a variety of insect pests (**Rossaro *et al*., 2018**). Sulfoxaflor is regarded as an alternative to, and even better than, some neonicotinoids because of their high potency and little cross-resistance with other pesticides (**Babcock *et al*., 2011**). The European Food Safety Authority reviewed the hazards of sulfoxaflor to humans, the environment, and non-target bodies when used as a pesticide in 2014, and reassessed the risks its in 2019 (https://www.greenfacts.org/en/sulfaxoflor-pesticide-bees/l-2/index.htm#0). Sulfoxaflor is somewhat poisonous to humans, birds, and most aquatic animals, but it is extremely dangerous to honeybees and bumblebees (**Al Naggar and Paxton, 2021; Chakrabarti *et al*., 2020**).

**Coragen 20% SC** is a novel insecticide that comprises the active component chlorantraniliprole, which has been licensed for use against insects in a variety of crops (**Kar *et al*, 2013**; **Bacca *et al*, 2021**). According to **Noha and Meligi (2019)**, toxicity coragen produced biochemical and histological modifications in various critical organs in male albino rats.

Spinetoram (**Radiant 12% SC**) is an insecticide that belongs to the spinosyn category and has a longer residual activity. Spinosad has a distinct method of action, working swiftly on the nervous system of insects by touch and ingestion (**Thompson *et al*. 2000**). Spinosyns are a novel class of insecticides with high activity and little environmental impact. Spinosyns have a distinct mode of action that involves the breakdown of nicotinic acetylcholine receptors (**Kirst, 2010**). Spinetoram is chemically related to spinosad, a pesticide with a proven track record of safety that may be used in organic farming. Spinetoram is particularly efficient against targeting insects at a low application rate while leaving beneficial insects alone. It works by consistently stimulating insect nicotinic acetylcholine receptors (**Anonymous, 2014**).

**Fipronil** is a potent pesticide that belongs to the novel phenylpyrazole pesticide class. It operates on the target organism by inhibiting GABA-gated chloride channels and glutamate-gated chloride (GluCl) channels (**Raymond, *et al*. 2005**). In laboratory circumstances, fipronil toxicity against land snail *Eobania vermiculata* was higher than that of land snails *Theba pisana* and *Helicella vestalis* (**Eshra, *et al*, 2016**). It was the most effective against hatchability percentage in the adult snails of ***Eobania vermiculata***, compared with untreated snails (**Hussein and Sabry, 2019**).

As a result, this study aims to assess the palatability of some plants leaves by Chocolate-band snail, ***E. vermiculata*** using two methods free-choice and no-choice, *in vitro*. Also, this labor tries to appraise the molluscicidal ability of five insecticides the Tetramic acid, Sulfoximine, Anthranilic diamide, Spinosad, and Phenylpyrazole classes against land snails, ***E. vermiculata***. Additionally, the effects of sublethal concentrations of these pesticides applied as leaf dipping on the activity of five essential enzymes in these land snails were investigated.

## Materials and Methods

### 1. Examined terrestrial molluscs

Adults’ snails of ***Eobania vermiculata*** were gathered by hand from Citrus limon trees during midden March 2022 from Buhaira Governorate. Animals were placed in transparent sacks and moved to the research center of the Zoology Agricultural and Nematology Department, Faculty of Agriculture, Al-Azhar University to be washed with freshwater (**Godan, 1983; Badawey, 2002**).

The snails were kept in a glass cage (60×40×30) cm filled up with mixed sterilized soil (clay: sand) roughly (1:1) with reasonable dampness and shut with a muslin material to avoid snails escaping from it. They fed on lettuce leaves for 15 days under lab conditions at RH 60%±5 and 22±1 ºC (**Shetaia, 2005; Bashandy and Raddy, 2021)**.

### 2. Tested plants

The foliage of six tested plants related to two families is displayed in Table (1)

**Table (1):** The foliage of tested plants for trial.

| **Table (1): The foliage of tested plants for trial** |  |  |
| --- | --- | --- |
| **family** | **Common name** | **Scientific name** |
| Asteraceae | chicory | *Cichorium cicorea Dumort.* |
|  | milky tassel | *Sonchus ciliatus Lam.* |
|  | Cos lettuce | *Lactuca scariola var. sativa* |
| Brassicaceae | London rocket | *Arabis charbonnelii* |
|  | Komatsuna | *Brassica rapa* |
|  | Cabbage | *Brassica oleracea* |

### 3. Daily food preferences of land snails by two methods

#### 3.1. no-choice feeding for terrestrial snails

Three plastic cups (10 depth × 14 diameters) filled with blend soil with 5cm depth, and 60% moisture were used for animals. Each cup had ten animals with three replicates. A well-known weight of plant leaves was given daily to animals. The consumption food of foodstuff was accounted for daily by snails and refilled again for five days. (**Eshra, 1997; Al-Akraa, *et al*., 2010 and Mohamed, 2016)**.

#### 3.2. Free choice feeding for land snails

Cabbage, Cos lettuce, Komatsuna, chicory, milky tassel, and London rocket leaves were used as test plants. Three wooden boxes (50×40×20cm) held mixed soil with a 10cm depth and 60% wetness. Each box included 30 animals, which were placed in the center of the box, and known weights of fresh leaf samples from each plant were placed around the snails on the box’s sides. To eliminate partiality for a certain place, the food ingredients and their sides were changed regularly (**Mohamed, Ghade, 2016**). After being weighed, the leaf samples were replaced every day. For five days, the adjusted weight losses caused by snail eating were estimated.

### 4. Pesticides utilized

Five pesticides in Table (**2**) were evaluated against adult snails of chocolate banded snail, ***E. vermiculata*** under lab conditions RH 60±5 and 22±1 ºC.

| N. | Common name | Trade name | Chemical group | Rate of application |
| --- | --- | --- | --- | --- |
| 1 | Spirotetramat | Movento 10% SC | Tetramic acid | 40 ml/100L |
| 2 | Sulfoxaflor | Transform 50% WG | Sulfoximine | 125 g/200L |
| 3 | Chlorantraniliprole | Coragen 20% SC | Anthranilic diamide | 60 ml/200L |
| 4 | Spinetoram | Radiant 12% SC | Spinosad | 10, 15 and 20 ml/feddan |
| 5 | Fipronil | Fipronil 5% | Phenylpyrazole | 500 ml/100L |

Concentrations were prepared based on the recommended field concentration. Seven concentrations of each pesticide were prepared. An appropriate size of lettuce leaf was dipped in each concentration and left for a minute, then taken out and left to dry, then served to snails. Ten animals were placed in each plastic box (15 x 10 cm) with three replicates. The cans are covered with a plastic lid with holes for ventilation. The plastic cans were checked daily for a month. Treated When needed, treated lettuce leaves were changed with untreated animals were sprayed with water to provide appropriate humidity. Dead animals were recorded and removed immediately.

### 3. Biochemical studies

#### 3.1. Preparation of the Sample

After seven days of treatment, the soft tissues of *E. vermiculata* and snails were removed from the shell and homogenized for one minute in 10 volumes (W/V) of 0.1 M phosphate buffer PH

7.4 using a glass homogenizer. The homogenates were centrifuged for 20 minutes at 1000xg in a cooling centrifuge (5417R) set to 4°C. The supernatants were kept in a deep freezer till used to determine the activities of alanine amino transaminase (ALT), aspartate amino transaminase (AST), lactate dehydrogenase (LDH), total protein (TP), and total lipid (TL). The supernatant was employed as an enzyme substrate (**Mourad and Zedan, 1996; Laila and Genena, 2011**).

#### 3.2. Determination of biochemical activities

The activities of aspartate aminotransferase (AST) and alanine aminotransferase (ALT) were assessed according to **Reitman and Frankel (1957)**. Lactate dehydrogenase (LDH) was assayed using the colorimetric method described by **Cabaud and Wroblewski (1958)**. Total proteins were assessed according to **Bradford (1976)**, and Total lipids were estimated according to **Knight *et al*. (1972)**. After this time, death percentages were computed and adjusted for natural mortality using Ldp line Software; **Bakr (2005)** to do probit analysis using **Finney (1971) and** according to **Duncan (1955) and Abbott’s method (1925)**.

## RESULTS

### 1. The current study used two approaches to determine which plant leaves were favored by land snails, *E. vermiculata*

#### 1.1. No-choice feeding for land snails

Data in Table (**3**) indicated that the average consumption weight of leaves of **Cos lettuce** was the best susceptibility with an average number of 4.769g for *E. vermiculata*. But cabbage, **Komatsuna, and milky tassel** recorded an average number of 3.961, 3.264, and 3.454g after 5 days, respectively. On the contrary, the **London rocket** was moderately palatable with an average consumption of 2.536 and chicory had the lowest palatable with an average consumption of 1.709g.

**Table (3):** *In vitro*, the no-choice consuming rate of some fresh plant foliage for the land snail, *Eobania vermiculata*.

| Plants |  | Cabbage | Cos lettuce | Komatsuna | chicory | milky tassel | London rocket |
| --- | --- | --- | --- | --- | --- | --- | --- |
| consuming rate/day | 1<sup>st</sup> | 3.858 | 4.647 | 3.091 | 1.704 | 3.949 | 2.653 |
|  | 2<sup>nd</sup> | 3.873 | 4.869 | 3.339 | 1.745 | 3.570 | 2.753 |
|  | 3<sup>rd</sup> | 4.453 | 4.548 | 3.387 | 1.661 | 3.259 | 2.436 |
|  | 4<sup>Th</sup> | 3.748 | 4.911 | 3.317 | 1.677 | 3.322 | 2.403 |
|  | 5<sup>Th</sup> | 3.873 | 4.871 | 3.185 | 1.759 | 3.168 | 2.436 |
| Mean |  | 3.961<sup>b</sup> | 4.769<sup>a</sup> | 3.264<sup>c</sup> | 1.709<sup>e</sup> | 3.454<sup>c</sup> | 2.536<sup>d</sup> |
| *Duncan's multiple range test shows that means with the same letter are not substantially different ($p \geq 0.05$). LSD 0.05 = 0.264* |  |  |  |  |  |  |  |

#### 1.2. Free choice feeding for land snails

According to the statistical analysis of the data in Table (**4**), there was a significant difference in average weight consumption for land snails (***E. vermiculata***). **Cos lettuce** and **cabbage** were the most prevalent with 3.828 and 3.392g eaten, respectively. **Chicory** also had a similar consumption rate to prior plants, with an average consumption rate of 3.625g. Furthermore, there is no statistically significant difference in the average intake of weight leaves of **milky tassel** and **London rocket** for land snails. On the other hand, the animal devoured the least amount of **Komatsuna** (1.606g) and had the lowest acceptance by snails for 5 days.

**Table (4):**
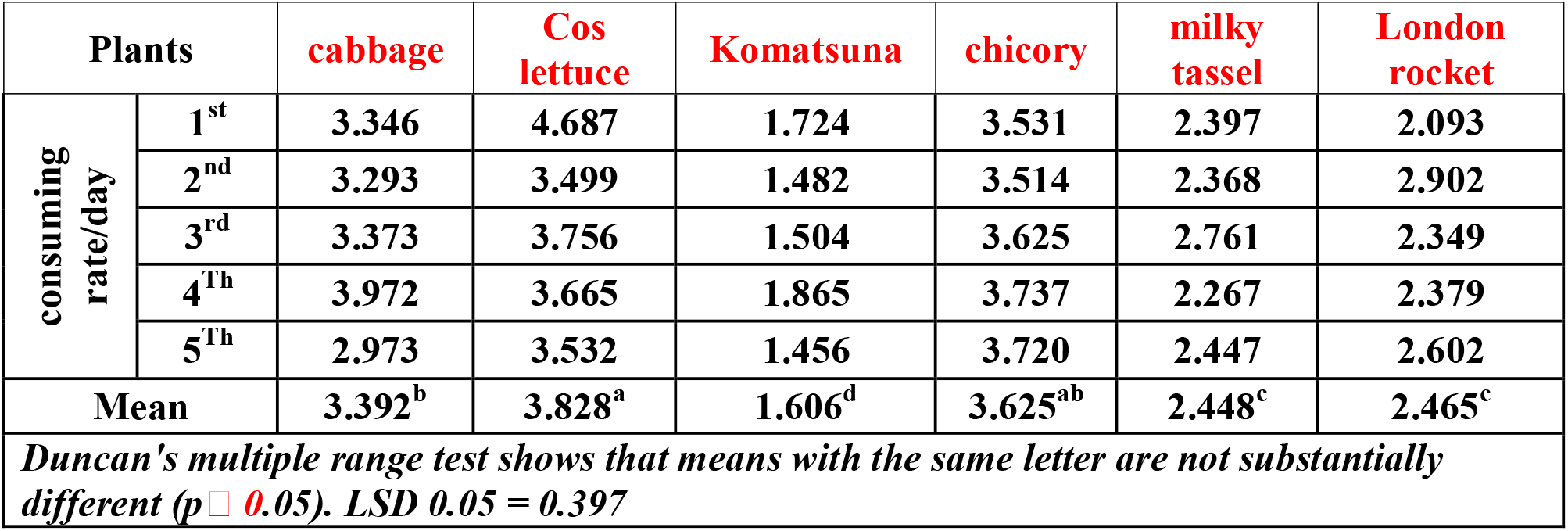
*In vitro*, the choice consuming rate of some fresh plant foliage for the land snail, *Eobania vermiculata*.

### 2. Efficacy of five pesticides applied as toxic leaf dipping against the brown land snail, *E. vermiculata*, under laboratory conditions

The molluscicidal activity of five insecticides applied as toxic disc lettuce leaf against the brown garden snail, *E. vermiculata* is shown in table (1). Data indicated the toxic effect of all tested insecticides leaf dipping with mortality percentage increasing with an increase in concentration and the period of exposure. From Table (**5**) and Fig. (1,2) it was evident that the tested pesticides Spirotetramat, Sulfoxaflor, Chlorantraniliprole, and Spinetoram showed no lethal effect against the land snail, *E. vermiculata* in the first six days after exposure.

**Table (5).**
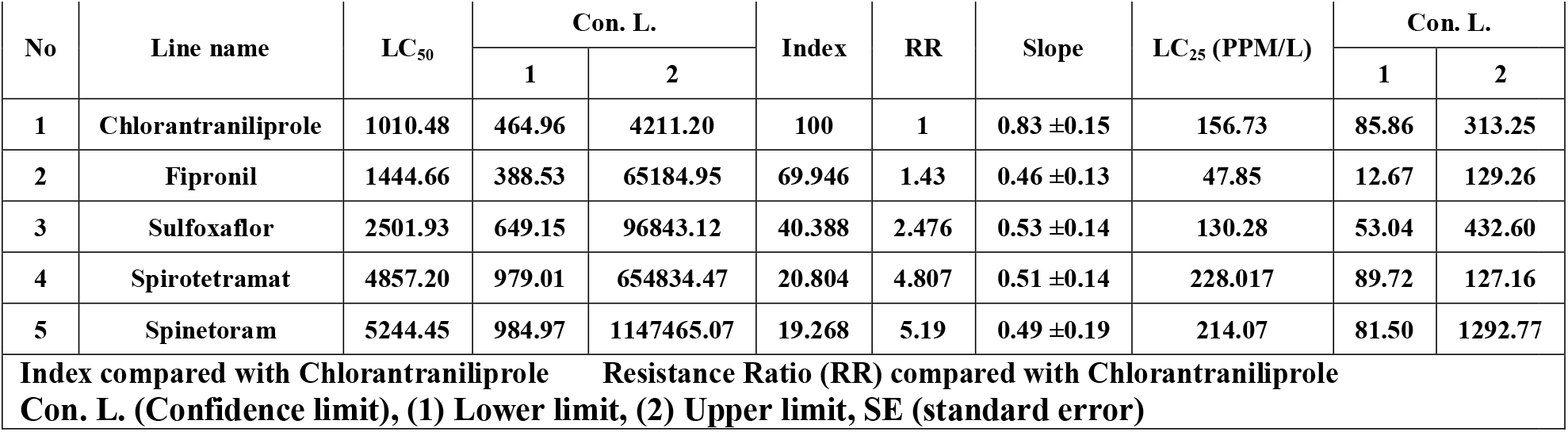
Molluscicidal activity (LC_50_ and LC_25_) of five insecticides against *E. vermiculata* using leaf dipping technique *in vitro*.

**Fig. (1):**
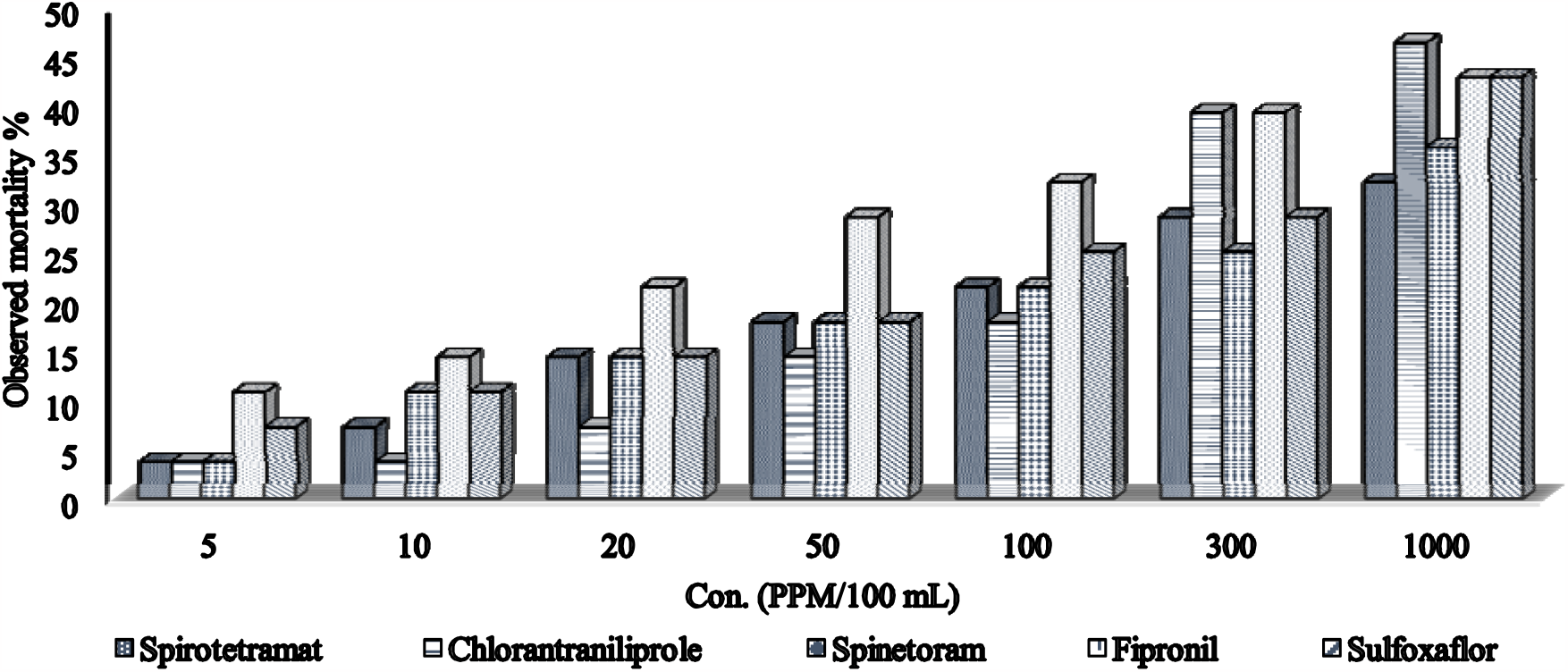
Percentage mortality of land snail, *E. vermiculata* by some insecticides for one month under laboratory conditions.

**Fig. (2):**
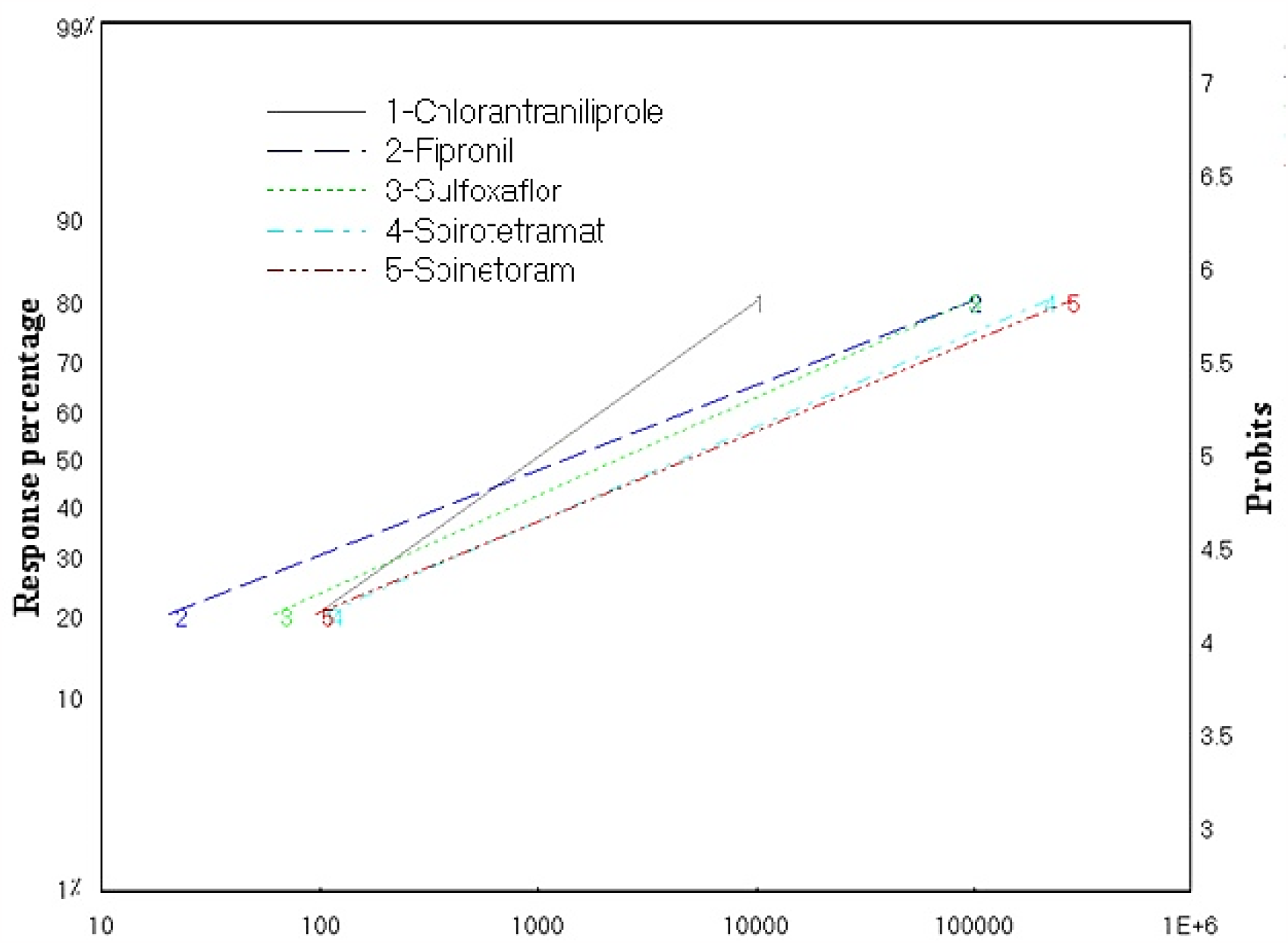
Probit regression lines representing the effect of insecticides leaf dipping against terrestrial snail, *E. vermiculata*.

However, after three days, the concentration of 5 ppm of sulfoxaflor and fipronil exhibited a mortality percentage of 7.142% and 10.714% for the brown land snail, E. vermiculata. While other insecticides displayed a low death rate in the tested animals at the same concentration.

After one week, the mortality percentage increased gradually for the tested insecticides. whilst after one month of exposure to 1000 ppm, Chlorantraniliprole significantly overwhelmed other pesticides and showed mortality of 46.43% with LC_50_ 1010.48 ppm/100ml. However, as the time elapsed to thirty days, all tested pesticides except Sulfoxaflor and Fipronil showed a gradual increase in the cumulative mortality percentage and exhibited a mortality percentage (42.86%) against *E. vermiculata* with lethal concentrations of 2501.93 and 1444.66 ppm/100 ml, respectively. Furthermore, in the final period trial, Spinetoram and Spirotetramat caused a lower percentage mortality of 35.71% and 32.14% with LC_50_ values of 5244.45 and 4857.20 ppm/100 ml, respectively. The untreated snails did not record any dead animals during the experiment.

In summary, the tested pesticides can be arranged in descending order according to their mortality percentages as follows: Chlorantraniliprole > Sulfoxaflor > Fipronil > Spinetoram > Spirotetramat.

### 3. Effects of five pesticides on (AST), (ALT), (TP), and (TL) in *E. vermiculata*

The obtained data in Table (6) show the effect of five pesticides mentioned perversely on (AST), (ALT), (TP), and (TL) in the land snail *E. vermiculata* after different periods.

**Table (6):** Effect LC_25_ of five different pesticides on enzymatic levels in the land snail *E. vermiculata* tissues exposed to 5ppm for 96 hrs.

|  |  |  |  |  |  |  |
| --- | --- | --- | --- | --- | --- | --- |
| Con. (ppm/L) per pesticides | ALT (U/L) | AST (U/L) | Total protein (g/dl) | lipid profile |  | LDH (U/L) |
|  |  |  |  | Total cholesterol (mg/dl) | Triglycerides (mg/dl) |  |
| Spirotetramat | 43<sup>c</sup> | 73<sup>e</sup> | 0.2<sup>a</sup> | 33<sup>a</sup> | 798<sup>c</sup> | 2<sup>a</sup> |
| Chlorantraniliprole | 38<sup>d</sup> | 164<sup>d</sup> | 0.3<sup>a</sup> | 1<sup>c</sup> | 271<sup>d</sup> | 1<sup>a</sup> |
| Spinetoram | 13<sup>e</sup> | 32<sup>f</sup> | 0.3<sup>a</sup> | 1<sup>c</sup> | 4728<sup>a</sup> | 1<sup>a</sup> |
| Fipronil 5% | 120<sup>a</sup> | 333<sup>c</sup> | 0.3<sup>a</sup> | 1<sup>c</sup> | 4<sup>f</sup> | 1<sup>a</sup> |
| Sulfoxaflor | 108<sup>b</sup> | 407<sup>b</sup> | 0.3<sup>a</sup> | 1<sup>c</sup> | 2064<sup>b</sup> | 2<sup>a</sup> |
| untreated | 37<sup>d</sup> | 448<sup>a</sup> | 0.2<sup>a</sup> | 3<sup>b</sup> | 6<sup>e</sup> | 2<sup>a</sup> |
| LSD<sub>0.05</sub> | 1.97 | 1.78 | 0.178 | 1.03 | 1.78 | 1.26 |
| Means with the same letter are not significantly different (p>0.05) according to Duncan's multiply range test.<br>ALT (alanine transaminase), AST (aspartate aminotransferase), and LDH (Lactate dehydrogenase).<br>U/L (units per liter), g/dL (grams per deciliter), mg/dL (milligrams per deciliter), |  |  |  |  |  |  |

#### 2.1. Effect on alanine aminotransferase (ALT)

When compared to other pesticides and controls, data showed that Fipronil and Sulfoxaflor increased ALT activity after seven days with 120 u/l and 108 u/l, respectively. The lowest ALT value achieved with Spinetoram after 7 days was 13 u/l. Spirotetramat and Chlorantraniliprole, on the other hand, achieved convergent outcomes with control levels of 43 and 38 u/l, respectively.

#### 1.2. Effect on aspartate aminotransferase (AST)

Radiant 12% and Spirotetramat substantially decreased AST activity following exposure days with 32 u/l and 73 u/l, respectively. Pesticides such as Sulfoxaflor, Fipronil 5%, and Chlorantraniliprole, on the other hand, began to raise the level of the enzyme after 7 days of treatment with 407 u/l, 333 u/l, and 164 u/l, respectively. In comparison to untreated animals throughout the same period, these results were the least effective on the activity of the enzyme AST.

#### 1.3. Effect on total proteins (TP)

The data demonstrated that there was no significant difference in total protein levels following testing between pesticides and controls.

#### 1.4. Effect on total lipids (TL)

Table (**6**) shows that when the four tested chemicals (Spinetoram, Sulfoxaflor, Spirotetramat, and Chlorantraniliprole) were administered to *E. vermiculata*, they raised the amounts of Triglycerides (4728, 2064, 798, and 271 mg/dl). Fipronil reduced the amounts of Triglycerides (4 mg/dl) in total lipids. All treatments and the comparator control had substantial changes. Seven days of Spirotetramat exposure resulted in the largest rise in Total cholesterol of lipid profile 33 mg/dl when compared to the control (3 mg/dl) and other pesticides (1 mg/dl). On total cholesterol, there is no significant difference between Chlorantraniliprole, Spinetoram, Fipronil, and Sulfoxaflor. The data demonstrated that there was no significant difference between pesticides and controls in the levels of LDH in the lipid profile after the exposure period.

## DISCUSSION

### 4. The current study used two approaches to determine which plant leaves were favored by land snails, *E. vermiculata*

**Nakhla and Tadros (1995)** discovered a strong preference for banana plants in E. vermiculata, orange trees in *H. vestalis*, the weed *Medicago polymorpha* in *T. pisana*, and orange trees in *Rumina decollate*. **Abd El-Hak (1997)** found that land snails, specifically *Monacha* sp. and *Eabania* sp., favored new lettuce leaves, followed by peas and cabbage, while garden rocket leaves were less preferred. **Arafa (1997)** reported the mean daily consumption (mg/snail) of *Eobania* sp. over 7 days, with lettuce, sweet peas, cabbage, and nursery rocket being consumed at rates of 1.1, 1.2, 1.4, 1.3, 1.5, 1.4, and 1.9 mg/snail, respectively. **Eshra (1997)** concluded that lettuce leaves were the most preferred, followed by cabbage leaves, while the fruits of carrot and squash were the least favored. **Shoeib (1997)** observed that *E. vermiculata* consumed more Dahlia leaves compared to cabbage and lettuce. On the other hand, **Mahrous, *et al*. (2002)** found that *Monacha cartusiana* snails preferred cabbage and lettuce in larger quantities, while pepper, pea, and tomato attracted fewer snails. **Mohamed-Ghada (2004; 2016)** demonstrated that land snails showed a preference for cabbage and lettuce over other plants. **Asran, *et al*. (2016)** revealed that E. vermiculata preferred lettuce, followed by squash, carrots, and potatoes. **Bashandy (2018)** observed that cabbage was the most palatable, with an average consumption rate of 0.552g, while London rocket and Snow thistle had the lowest consumption rates for the march slug, *Deroceras leave*, at 0.244g and 0.215g, respectively.

### 2. Efficacy of five pesticides applied as toxic against the brown land snail, *in Vitro*

**Pesticides and Authority (2009)** found that over the first 28 days, death in oral and contact toxicity techniques by spirotetramat with predatory mites were exposed varied from 90.5-100%. The highest test concentration, however, dramatically reduced mortality in gamasid mites (*Hypoaspis aculeifer*) on day 21 and set the LC_50_ value to >1000 mg metabolite/kg dry soil in comparison to the control. **Garcerá *et al*. (2013)** showed that spirotetramat is efficient in lowering *Aonidiella aurantii* stages in the laboratory. Females in the first and second instars of *A. aurantii* were 10- and 32-fold more sensitive to spirotetramat (LC_50_ =0.1-0.2 ppm) than females in the early third (LC_50_ = 1.5 ppm) and late third (LC_**50**_ = 5.3 ppm). Spirotetramat would also diminish fecundity by 89%. **Al-kazafy and Mona** (**2022**) envaulted that spirotetramat on land snails, *E. vermiculata* by conventional and nanoformulations that achieved 100% and 53% mortality with LC_50_ (7.7 and 35%) with increasing concentrations, respectively.

**Kumar *et al*. (2013)** investigated the harmful impact of subacute oral administration of Coragen at a dosage of 1000 mg/kg body weight after 14 and 21 days of exposure on several hematological parameters in rats. According to **Deleva and Harizanova (2014)**, the pesticide chlorantraniliprole (Coragen® 20 SC) induced 100% larval mortality of *Tuta absoluta* (Meyrikc), a significant tomato pest, in lab conditions. Coragen (chlorantraniliprole) produced alterations in the pathological parameters of the sub-acute and sub-chronic liver, kidneys, and protein profile on Albino Rat, resulting in an increase in AST and ALT activity, urea, and creatinine concentrations, according to **Abdel-Mobdy *et al*. (2017)**. In the fish *Perca fluviatilis*, **Ponepal *et al*. (2023)** discovered that Coragen 20 SC produced behavioral changes, a decrease in respiratory rate and oxygen consumption, an increase in blood glucose levels, and a decrease in the number of erythrocytes and leukocytes. At the applied concentrations (0.0125, 0.025, and 0.05 ml/L), its toxicity on fish *Triticum aestivum* was poor to moderate phytotoxicity for the evaluated parameters.

**Eshra, *et al*., (2016)** evaluated fipronil as molluscicidal belonging to phenyl pryazole against *Eobania vermiculata* under laboratory conditions using a leaf-dipping bioassay method. The result revealed that the LC_50_ value (after four days) was 0.014% for fipronil against land snail. **Mona and Al-Kazafy (2019)** showed that the recommended field rate of Fipronil was very effective against the eggs of *E. vermiculata* at 22.7% compared with 96.3% in control.

**El-Bassouiny, *et al*., (2022)** demonstrated that Spiroetramat (Movento) achieved a 24 h-LC_50_ of 12.05 ppm against cotton bollworm, *Earias insulanaare*, and caused a significant change in the activities of transaminases enzymes (AST and ALT), phenoloxidase, and acetylcholinesterase. Total protein and lipids had the highest significant decrease.

Metabolization of sulfoxaflor in both animals and plants occurs through oxidative cleavage at the methyl(oxo) sulfanylidene cyanamide moiety, and it can proceed through glucuronidation to form conjugates (**Pfeil, *et al*. 2011**). According to **Gore *et al*. (2013)**, the LC_50_ values for sulfoxaflor against cotton aphids were 1.01 to 5.85 ppm and 0.92-4.13 ppm at 48 and 72 hours, respectively. After 15 days, **Das *et al*. (2019)** discovered that Sulfoxaflor 50% WG was a more effective dosage against tea mosquito bug (*Helopeltis theivora*) and Green Fly (*Empoasca flavescens*) than check pesticides Thiamethoxam 25% WG and Clothianidin 50% WDG. Sulfoxaflor, as demonstrated by Li et al., 2021, is a new sulfoximine pesticide that is commonly used to treat crop pests. They studied the effects of sulfoxaflor on larval and newly emerged worker honeybees. It was toxic to adult bees and caused significant changes in antioxidative (SOD, CAT), lipid peroxidation (POD, LPO, MDA), detoxification (GST, GR, GSH), and signal transduction-related (AChE, ACh) enzymes or products in both larvae and adult honeybees in the laboratory over 96 hours.

According to **Al Naggar and Paxton (2021)** and **Chakrabarti *et al*. (2020)**, sulfoxaflor is moderately toxic to mammals and birds and slightly toxic to most aquatic species, but it poses a high risk to honeybees and bumblebees when they come into contact with spray droplets shortly after application. Some studies have reported the high toxicity of sulfoxaflor to bees. Sulfoxaflor is registered as having a contact acute LD_50_ of 0.379 μg/bee and an oral acute LD_50_ of 0.146 μg/bee for honeybees (*Apis* spp.) in the PPDB (Pesticide Properties Data Base, http://sitem.herts.ac.uk/aeru/ppdb/en/Reports/1669.htm).

### 3. Effects of five pesticides on some biochemical parameters in *E. vermiculata*

Pesticides produce biochemical impairment and lesions of tissues and cellular processes increasing the activity of enzymes multiplied hundreds of times ordinary such as (AST), (ALT), (ALP), and (LDH) in land snails, *Monacha cantiana*, and *Theba pisana*, due to cell damage in their many organs (**Felix Wróblewski, *et al*., 1956; Radwan, *et al*., 1992; Farkas, *et al*., 2004; Arceci, *et al*., 2006; Mahmoud, 2006; Celik, *et al*., 2009; Ghouri, *et al*., 2010 and El-Gohary and Genena, 2011 and Khalil, 2016**). Also, **Bakry *et al*. (2013)** found that exposing *Bulinus truncatus* to sublethal amounts of glyphosate for two weeks elevated the activities of lipid peroxidase (LP). Additionally, the tested chemical compounds on *E. vermiculata* changed a number of biological targets, which may lead to serious negative effects on snails’ metabolism and cells (**Mahal, *et al*., 2015**). According to **Abdelmonem (2016)**, methomyl is hazardous to non-target creatures such as land snails. Lannate had an LD_50_ of (102.32 μg/snail) and inhibited AChE more in the brain ganglia than in the foot muscle. Except ALP, there was a considerable elevation of hemolymph enzymes in snails exposed to 21.32 and 53.30 μg/snail for 48 hours via contact poisoning on *Eobania vermiculata*. As well, **Esam (2023)** showed that chemical compounds treatment of *E. vermiculata* decreased the activities of ALT, AST, and TP with mean values less than the control, while a mixture of chemicals treatment increased ALT, AST activities, and TP with mean values greater than the control.

## Conclusion

In conclusion, the results showed that land snails displayed a preference for both Cos lettuce and cabbage leaves in both the “no-choice feeding” and “free-choice feeding” methods. The tested pesticides can be arranged in descending according to their mortality percentages as follows: Chlorantraniliprole > Sulfoxaflor > Fipronil > Spinetoram > Spirotetramat. Also, Fipronil and Sulfoxaflor increased the activity of ALT with 120 u/l and 108 u/l and Sulfoxaflor, Fipronil, and Chlorantraniliprole began to increase the level of the enzyme AST after 7 days of treatment with 407 u/l, 333 u/l, and 164 u/l, respectively. Spinetoram caused the lowest value of ALT and AST with 13 u/l and 32 u/l, respectively. Whereas Fipronil caused a reduction of Triglycerides (4 mg/dl) of TL. But there is no significant difference between pesticides and control affected the total protein, (Total cholesterol and LDH) of lipid profile levels after the exposure period. Therefore, these substances could be useful for managing land snails. The most likely route of action of these chemicals in land snails still needs more research.

